# Forest dieback in drinking water protection areas – a hidden threat to water quality

**DOI:** 10.1101/2024.08.07.606951

**Authors:** Carolin Winter, Sarina Müller, Teja Kattenborn, Kerstin Stahl, Kathrin Szillat, Markus Weiler, Florian Schnabel

## Abstract

For centuries, forests have been considered a natural safeguard for drinking water quality. We challenge this view in light of the rising frequency of climate extremes, such as droughts coinciding with high temperatures, which have caused unprecedented pulses of forest dieback globally. Drought-induced forest diebacks may jeopardize the crucial role of forests in protecting water quality, potentially even turning forests into sources of contamination. To underscore the critical importance of the topic, here we provide the first comprehensive assessment of forest cover, type, and dieback (assessed as canopy cover loss) across drinking Water Protection Areas (WPAs) in Germany, one of the countries hit most severely by the unprecedented Central European drought of 2018–2020. Our findings reveal a high forest cover of 43% in WPAs, from which a substantial amount of 5% canopy cover got lost within only three years as a direct or indirect consequence of the drought. Spruce-dominated forests, constituting 28% of all forests in WPAs, were particularly susceptible, but other dominant tree species also experienced anomalously high mortality rates. Combining this assessment with exemplary records of nitrate concentrations in the groundwater of WPAs revealed that forest dieback can significantly impair drinking quality. On average, nitrate concentrations more than doubled in WPAs with severe forest dieback, whereas nitrate concentrations did not significantly change in undisturbed WPAs. However, we also found pronounced differences between WPAs affected by forest dieback, underlining the need for further data and research to derive a generalizable understanding of the underlying mechanisms and controls. Based on this assessment, we deduce critical data and knowledge gaps essential to developing well-informed prediction, adaptation, and mitigation strategies. We call for interdisciplinary research addressing the hidden threat forest dieback poses for our drinking water resources.

## 1 INTRODUCTION

For centuries, forests in drinking Water Protection Areas (WPAs) have been conceived as a safeguard for drinking water quality. WPAs, located in the realm of drinking water abstraction facilities, are key-areas to ensure local or regional drinking water supply. The mere presence of semi-natural forests, hereafter ‘forests’, in WPAs already excludes intensive land use and pollution sources prevalent in agricultural, urban, or industrial areas (Hegg et al., 2004; Piffer et al., 2021). A very common contaminant in many forests is nitrate, a nutrient whose high concentrations threaten the health of aquatic ecosystems and human health if ending up in our drinking water (Bijay-Singh & Craswell, 2021; Elser, 2011; Vitousek et al., 1997; Ward et al., 2018). In forests, nitrate primarily stems from atmospheric nitrogen (N) deposition (Hegg et al., 2004; Winter et al., 2021). Nitrate is highly mobile and, in excess, rapidly leaches into the groundwater. Groundwater, in turn, is an invaluable source of drinking water that accounts for approximately half of the domestic water supply worldwide (UN, 2022). Besides their passive role in protecting drinking water quality, forests actively take up nutrients (e.g. nitrate from atmospheric deposition) and keep them in closely tied cycles, thereby buffering nutrient leaching into the groundwater (Figure 1A). Moreover, forest ecosystems enable the formation of humus-rich soil layers that enhance biological activity and nutrient absorption (Hegg et al., 2004). As a result, water in or from forested areas often maintains a high quality (Dudley & Stolton, 2003; Karr & Dudley, 1981; Mupepele & Dormann, 2017; Piffer et al., 2021).

**Figure 1.**
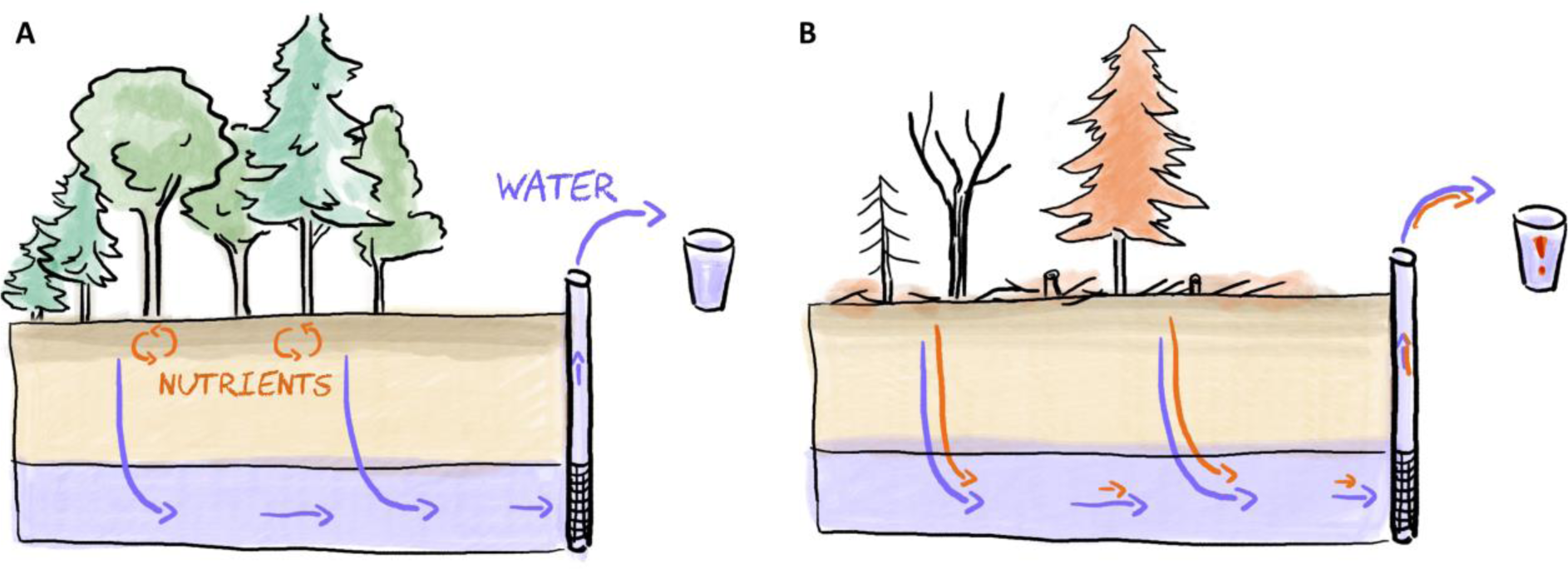
Schematic view on the role of forests as a safeguard of groundwater quality and related drinking water quality.

However, the increasing frequency, intensity, and duration of climate extremes severely threaten the vitality and survival of forests worldwide (Adams et al., 2009; Breshears et al., 2005; Cheng et al., 2024; Hartmann et al., 2022). Across Europe, Senf et al., (2020) found clear evidence for drought-induced excess tree mortality (i.e., tree mortality exceeding the long-term reference). Moreover, droughts interact with other disturbance factors such as pests and pathogens or storms, which jointly induced a significant increase in disturbance impacts across European forests since 1950 (Patacca et al., 2023). A striking example is the drought of 2018–2020, which set a new benchmark across Central Europe due to its intensity, duration, and high temperatures (Rakovec et al., 2022). The extremely dry and hot year of 2018 alone led to unprecedented rates of tree mortality (Schuldt et al., 2020). However, in response to the consecutive drought years of 2019 and 2020, tree mortality surged even further, driven by the cumulative stress these drought years exerted (Obladen et al., 2021; Schiefer et al., 2023; Schnabel et al., 2022). As a result, the German Forest Condition Survey reported that in 2023, only about every fifth tree was in good condition, and around 6.7% of trees died (BMEL, 2024). Worldwide, such drought-induced forest dieback events are increasing across forested biomes, even those previously not considered at risk (Hartmann et al., 2022).

The recent emergence of drought-induced forest dieback may jeopardize the crucial role of forests in protecting water quality and challenge the prevailing view of forests as a safeguard for drinking water quality. Different case studies have shown elevated nitrate concentrations in soil or surface water after forest dieback in areas with standing deadwood (Clow, 2010; Huber, 2005; Mikkelson, Bearup, et al., 2013; Mikkelson, Dickenson, et al., 2013) or after tree removal (Dahlgren & Driscoll, 1994; Kong et al., 2022). One reason is that dead trees do not take up nutrients anymore. Hence, the forests lose a critical part of their ability to buffer N inputs (Figure 1B). An additional lack of transpiration can promote a higher water availability for nitrate leaching (Adams et al., 2012; Clow, 2010). In addition, the decay of organic material, such as leaves, needles, twigs, and roots, releases nutrients that can be transported to the groundwater (Pearson et al., 1987). Mineralization and nitrification, accelerated by higher temperatures, higher light availability, and a shift in the soils’ C:N ratio, can release nutrients stored in humus-rich soils (Borken & Matzner, 2004; Göttlein et al., 2003; Kopáček et al., 2017). In summary, damaged forests cannot only lose their role in protecting drinking water resources but can also transform into an additional source of contamination.

Still, we lack a generalizable understanding of the dominant drivers and controls on drinking water quality in areas affected by forest dieback. While much research has focused on nutrient dynamics in forests, it has mainly been limited to single stands and concentrated on soil water (e.g., Adams et al., 2012; Huber, 2005; Mikkelson, Bearup, et al., 2013) or surface waters (e.g., Clow, 2010; Kong et al., 2022; Kopáček et al., 2017). Hence, it remains unclear if changes in the shallow water quality ultimately reach the groundwater, which is the most important source of drinking water in Germany and many other countries (Destatis, 2018; UN, 2022). Moreover, studies are missing that compare impacts on groundwater quality across different forest types, types of forest damage (e.g., drought, wind throw, insect infestations, wildfire or compound events of these), and management strategies (salvage cutting, leaving trees standing, active/passive reforestation, etc.). Such knowledge would be essential for understanding the mechanisms of how different types of forest dieback affect drinking water quality and developing effective prediction, adaptation, and mitigation strategies.

To close this gap, we argue that interdisciplinary research spanning water chemistry, hydrology, forestry, soil science and remote sensing is indispensable. We need an improved understanding of the interlinked processes related to forest dieback and hydrological transport below the root zone, as well as approaches that allow upscaling of these observations to scales relevant to management and decision-making. In support of our call for intensified research in this direction, here we first present the potential dimension of the issue, i.e., the first assessment on forest cover, forest type, and dominant tree species across all WPAs in Germany. Second, we assess forest dieback across all WPAs in Germany in response to the Central European drought of 2018–2020, one of the countries that were hit most severely by this unprecedented global-change type drought and its related forest dieback (Hartmann et al., 2022; Rakovec et al., 2022). In a third step, we zoom in and examine time series of groundwater nitrate concentrations in a subset of WPAs with and without severe forest dieback, demonstrating the potential effect of forest dieback on drinking water quality. Based on this assessment, we highlight future challenges (i.e., critical data and knowledge gaps) and present ways forward to address the hidden threat forest dieback poses for our drinking water resources.

## 2 The potential dimension: Forest cover and -type in German drinking water protection areas

Here, we present the first German-wide dataset of WPAs, compiled from all 16 Federal States’ designated protection areas that were available from each of the official state-geographical information systems (see data availability statement). While WPAs generally underlie restrictions to minimize the contamination risk, their definition slightly differs among the Federal States. Therefore, the resulting areas can represent slightly different legal protection states or even some areas without legal protection. Nevertheless, this dataset enabled us to compile the first assessment of forest cover and type in Germany’s WPAs. This assessment allowed us to create an overview of areas whose drinking water resources have, so far, been protected by forests but that might lose their protective role or even transform into an active source for drinking water contamination in the face of forest dieback (Figure 2).

**Figure 2.**
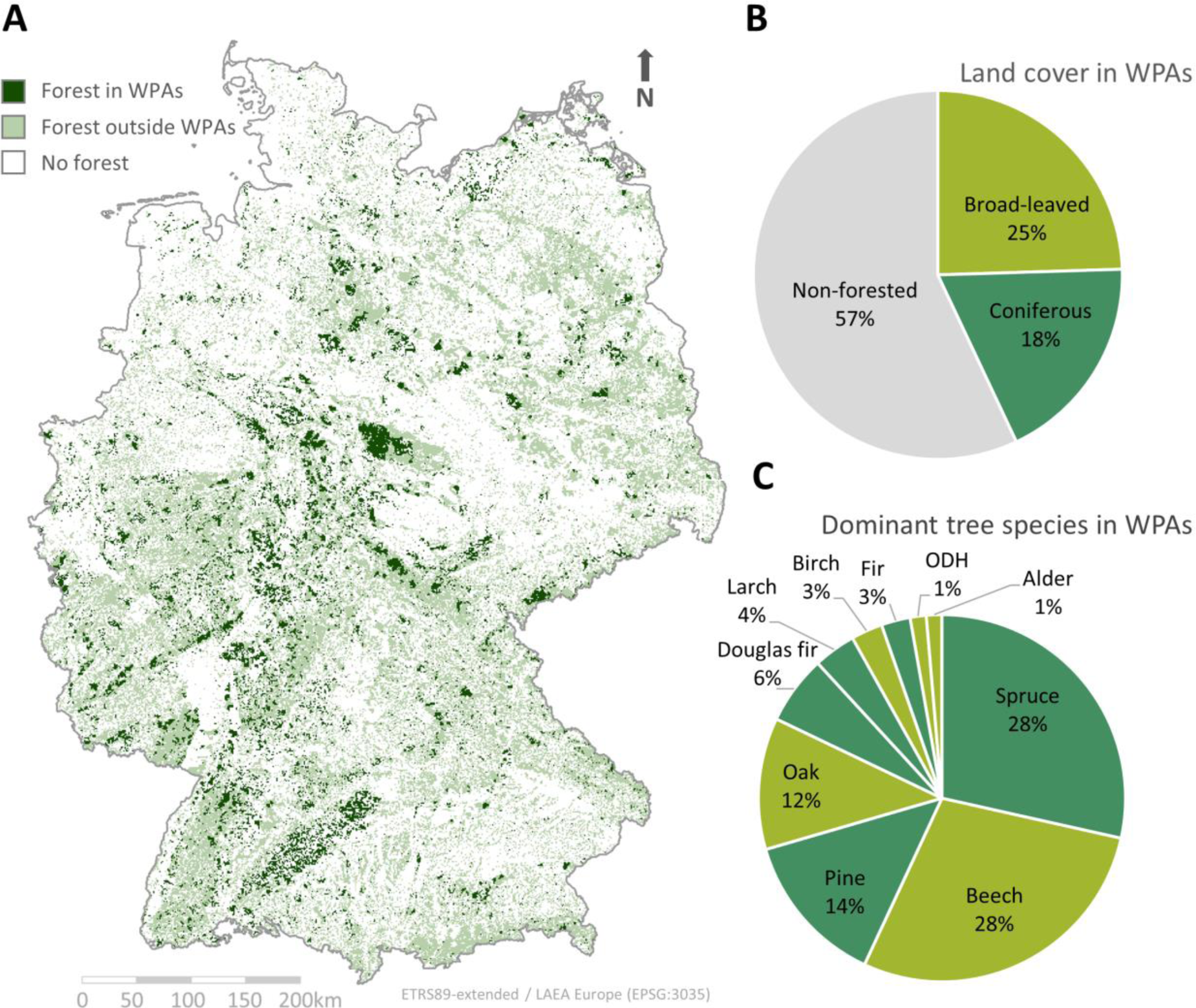
Overview on forest cover, forest type and dominant tree species in drinking water potection areas (WPAs) across Germany. A) A map depicting the spatial distribution of forests inside and outside of Germany’s WPAs, B) a chart depicting the relative area contribution of broad-leaved versus coniferous dominated forests, and C) the percentage of dominating tree species in Germany’s WPAs.

To assess the percentage of forest and forest type (i.e., coniferous versus broad-leaved forest), we used the Forest Type dataset provided by the Copernicus Land Monitoring Service at a 20m x 20m resolution (EEA, 2018). We further identified the dominant tree species growing across German WPAs, using the dataset provided by the Thünen Insitute, which is based on Sentinel-1 and Sentinel-2 data from 2017 and 2018 (Blickensdörfer et al., 2022).

Combined, these data sources show that approximately 43% of German WPAs are covered by forest (Figure 2). This is a disproportionally high coverage compared to 32% forest cover across the entire country. Overall, there are more broad-leaved than coniferous dominated forests, while dominant tree species groups are spruce (*Picea*) and beech (*Fagus*), followed by pine (*Pinus*) and oak (*Quercus*). Note that the percentages of forest type and dominant tree species cannot be set off against each other, as they relate to slightly different bases. To illustrate, even if broad-leaved beech would be dominant in one pixel – spruce, douglas fir, and fir could still make up >50% of it, making it a predominantly coniferous forest. In summary, a large share of WPAs in Germany is covered by forest that spans a range of different forest types and dominating tree species.

## 3 Ongoing damage: forest dieback in response to the drought of 2018–2020

Forest dieback, which we assessed here as canopy cover loss, strongly increased from 2018 onwards in German WPAs (Figure 3B). Hence, forest dieback in German WPAs followed similar trajectories as in the rest of Central Europe (Schuldt et al., 2020; Senf et al., 2020). By April 2021, approximately 5% of canopy cover had been lost within approximately three years, which is a similar percentage as the one reported by Thonfeld et al. (2022) for entire Germany. Out of these 5%, 92% were coniferous and 8% broad-leaved forests (Figure 3A). These 5% are a substantial loss, especially in WPAs that are critical areas for local and regional drinking water supply and considering that the rotation length of dominant tree species in Germany ranges from about 60–160 years.

**Figure 3.**
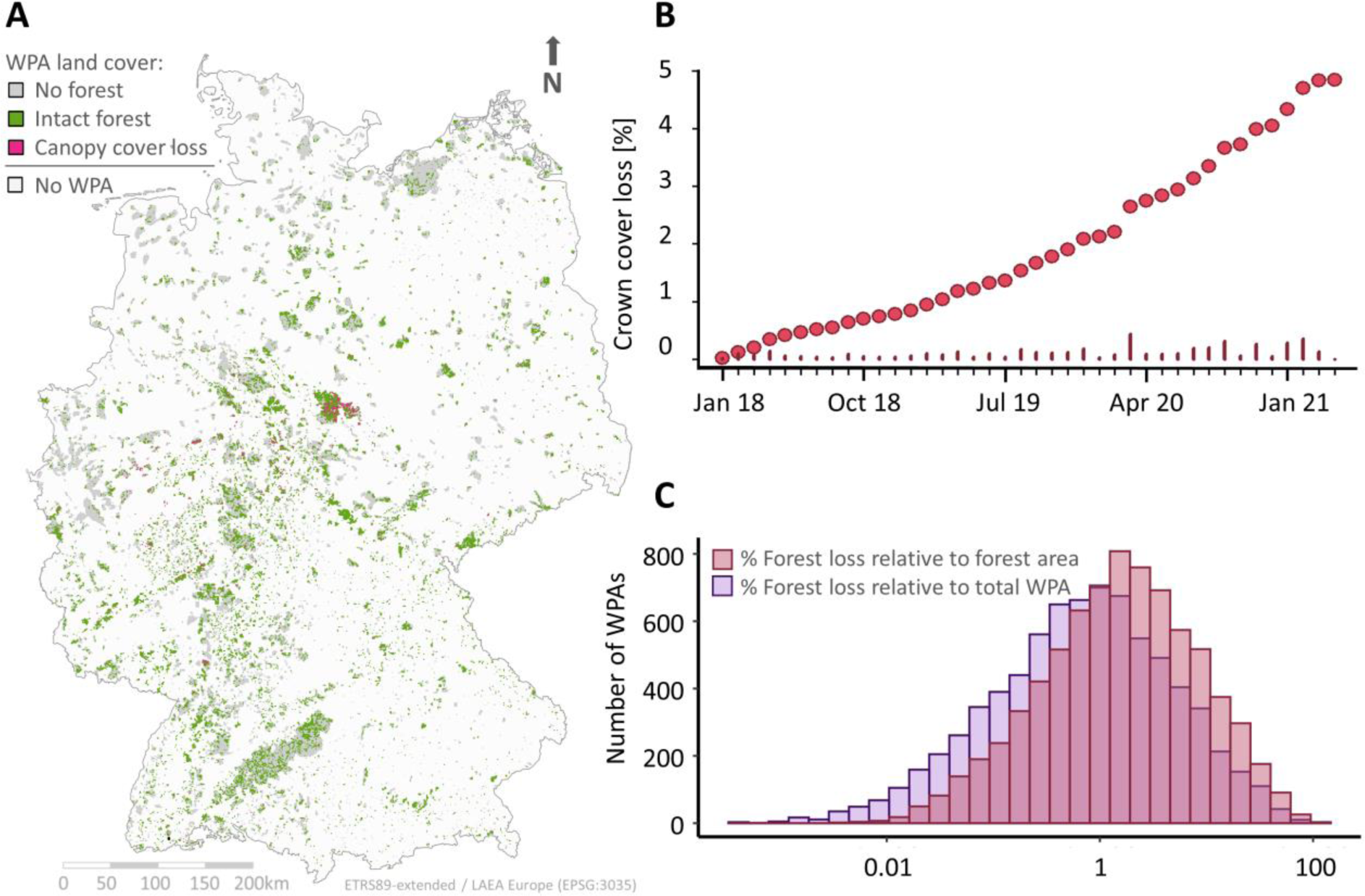
Canopy cover loss across German WPAs, with A) a map of intact forest and canopy cover loss between January 2018 and April 2020, B) the temporal resolution of canopy cover loss as absolute (lines) and cumulative (dots) percentage, and C) the percentage of canopy cover loss relative to the forested area of a WPA and relative to the entire area of the WPA.

Canopy cover loss occurred predominantly around Central Germany, where the drought was most severe (Hari et al., 2020; Rakovec et al., 2022). However, also the remaining 95% of ‘intact’ forests in WPAs are largely in an exceptionally poor condition after the drought of 2018–2020 (BMEL, 2024). Drought legacy effects likely further reduce the forests’ resistance to subsequent abiotic and biotic stress and disturbance agents, predisposing these forests to future forest dieback events (Anderegg et al., 2015; Obladen et al., 2021; Schnabel et al., 2022).

Specifically, we used canopy cover loss data (i.e., standing dead or removed trees) from Thonfeld et al. (2022) to assess the extent of forest dieback over WPAs in response to the drought of 2018–2020. This data is based on Sentinel-2 and Landsat-8 data with a spatial resolution of 10m x 10m and a monthly resolution starting in January 2018 and lasting until April 2020. Canopy cover loss was estimated via the anomaly of the disturbance index (DI) introduced by Healey et al. (2005) and identified using a fixed threshold for the difference between the 10^th^ percentile of the DI per month starting in 2018 and the median DI of 2017 used as the reference year with ‘normal’ climatic conditions.

Overall, we counted 317 WPAs with a substantial canopy cover loss >25% off the forested area, and 176 WPAs with a canopy cover loss >25% off the total WPA (Figure 3C). From these 317 and 176 WPAs, the majority were dominated by spruce (n=251 and n=160), followed by beech (n=34 and n=13), and other tree species like pine, douglas fir, fir and oak. These numbers show that coniferous and mainly spruce-dominated forests have been affected most by forest dieback, in line with former reports for the entire Germany (e.g., BMEL, 2024; Thonfeld et al., 2022). One reason is that spruce is often planted in monocultures and out of its natural distribution range, making it especially sensitive to climate extremes and insect infestations (Jactel et al., 2021). Besides the drought event of 2018–2020 (see, e.g. Obladen et al., 2021), spruce forests have been severely affected by wind throws from storms in the past decades, for example, by Lothar in 1999 or Kyrill in 2007, and by interactions between bark beetle outbreaks, warmer temperatures and drought (Biedermann et al., 2019; Patacca et al., 2023). Consequently, the high susceptibility of spruce to dieback, in conjunction with more than a quarter of all forests in WPAs in Germany being dominated by spruce (Figure 2), may pose a substantial threat to drinking water quality. Nevertheless, we also observed anomalously high mortality rates in other dominant tree species, such as beech and fir, in response to the drought of 2018–2020, which is in line with the German Forest Condition Survey (BMEL, 2024). The scale of these numbers raises the question of whether and which of the contemporary tree species in Germany can still contribute to safeguarding drinking water quality in the face of accelerating climate change.

## 4 Is drinking water quality affected by forest dieback?

To test whether we are able to elucidate the threat forest dieback might pose to drinking water quality, we analyzed nitrate concentrations in the groundwater of WPAs with severe canopy cover loss. To this end, we selected a subset of all WPAs with >90% pre-drought forest cover (to avoid confounding effects from agriculture or urban areas) and >25% canopy cover loss. As a reference, we selected 6 sites with >90% forest cover but <3% canopy cover loss that are located in similar regions as the sites with severe forest dieback (Figure 4). From the potential 114 WPAs with high canopy cover loss, only 13 sites had freely available nitrate concentration data provided by the Environmental Agencies of the Federal States of Germany, which were measured from wells or springs inside of the WPA and covered the time period from 2008–2021 or longer (Figure 4C; Table S1). These sites give an exemplary overview on local impacts of forest dieback on groundwater quality. However, due to their low number and clustering in Central Germany, where most forest dieback occurred, this assessment can only serve as a starting point for future investigations. Further details can be found in the Supplement (Table S1).

**Figure 4.**
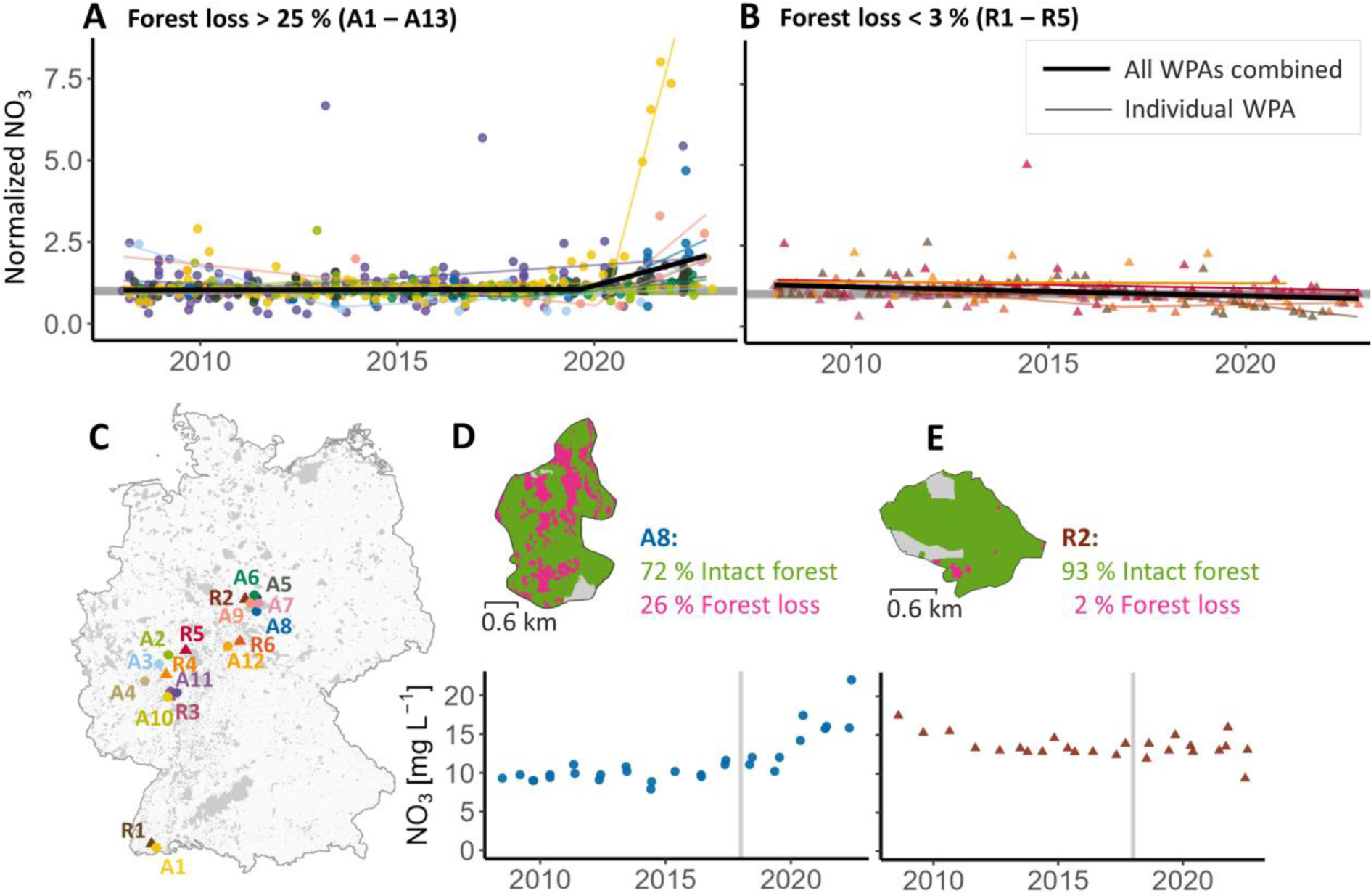
Normalized groundwater nitrate concentrations between 2008 and 2022 in forested sites that were A) affected by <25% canopy cover loss or B) reference sites with <3% canopy cover loss. Thin colored lines depict the area-specific linear models with or without a breakpoint. Thick black lines depict the linear models across all sites with >25% or <3% canopy cover loss. Panel C) depicts a map with the locations of the canopy cover loss and reference sites. Panels D) and E) show two exemplary sites, one with substantial canopy cover loss and a significant increase in nitrate concentration after 2018 (D), and the other a reference site with marginal canopy cover loss and no significant breakpoint after 2018 (E).

We found a pronounced increase in groundwater nitrate concentrations in response to forest dieback in more than half of the WPAs with severe canopy cover loss (>25%). Seven out of 13 sites showed a significant breakpoint, with increasing nitrate concentrations after 2017 (Figure 4A, D). In contrast, none of the reference sites had a breakpoint in that period (Figure 4B, E). The breakpoint analysis was conducted using the segmented package in R (Fasola et al., 2018; R Core Team, 2024) and applied to the deviation of nitrate concentrations from the site-specific median. Breakpoint analysis was conducted for each WPA individually and in a combined approach across all WPAs affected by severe forest dieback as well as across reference sites (Figure 4A, B). Combined, sites with severe canopy cover loss had a significant breakpoint in December 2019, after which nitrate concentrations started to increase drastically, while no significant breakpoint across all reference sites could be detected. In absolute numbers, median groundwater nitrate concentration across all sites between 2008 and 2017 was 5 mg L^-1^ (IQR: 2–8 mg L^-1^). After forest dieback in 2021 and 2022, nitrate concentrations across the affected sites more than doubled, reaching a median of 11 mg L^-1^ (IQR: 7–15 mg L^-1^), whereas nitrate concentrations in the reference sites remained at a median of 5 mg L^-1^ for both periods. We note that none of the sites has so far exceeded the threshold for nitrate concentrations in drinking water (50 mg L^-1^; WHO), but a further increase might change this (see our argumentation below). Moreover, groundwater from forested sites that has low nitrate concentrations is currently an important resource for diluting groundwater with high nitrate concentrations from agricultural sites (LfU, 2013), a role that may be increasingly lost under forest dieback.

The breakpoint for affected sites in the end of 2019 lies more than a year after the start of the 2018 drought. This delayed response can likely be explained by a mix of cumulative stress responses of the trees after two consecutive drought years (Schnabel et al., 2022) and the time it takes for hydrological nitrate transport to the groundwater. This hydrological transport, often described by water age or transit time distributions of water and nitrogen, can range from a few months to several decades (e.g., Jutglar et al., 2021; Osenbrück et al., 2006; Visser et al., 2013; Winter et al., 2021, 2023), suggesting that what we see is likely just the start of groundwater quality deterioration. Even if looking at the soil water alone, which should respond faster than groundwater, Huber (2005) already reported highest nitrate concentrations 5 years after forest dieback and lowest concentrations 17 years after the dieback (although with no measurement from year 8 to 15). Consequently, we expect that the groundwater nitrate concentrations we report will increase even further and that concentrations will remain at a high level for several years to decades to come.

Finally, pronounced site-specific differences in the timing and magnitude of change became apparent (Figure 4A). Six out of the 13 sites did not show a significant breakpoint, meaning that groundwater nitrate concentrations were either unaffected or the effect is delayed and not yet visible. One reason for these differences could be different hydrological flow paths and a related delay (see above). However, other abiotic or biotic factors, such as differences between forest or disturbance types, might also play a crucial role. For instance, spruce forests, which make up the majority of affected WPAs, are also the ones with the highest atmospheric N-deposition and nitrate leaching compared to other forests in Germany (Borken & Matzner, 2004), which may lead to an amplification of water quality deterioration. This underlines the need for further research on underlying drivers and controls of nitrate concentration changes after drought-induced forest dieback.

## 5 Future challenges

Using Germany as an example, we demonstrate that substantial forest dieback as a direct or indirect consequence of drought events can significantly deteriorate groundwater quality in WPAs. Our findings revealed a substantial 5% canopy cover loss across WPAs in Germany following the drought of 2018–2020. Spruce-dominated forests were particularly susceptible, but other dominant tree species also experienced anomalously high mortality rates. This indicates that contemporary tree species cannot guarantee vital forests which protect drinking water quality in the face of climate change. In over half of the observed WPAs, forest dieback significantly deteriorated groundwater quality, with average nitrate concentrations more than doubling during the observation period.

Although this is an alarming trend, pronounced differences among sites highlight the need for further research on the underlying drivers and controls of nitrate concentration changes following forest dieback. The situation assessed stands exemplary for many countries in the world with forest-reliant drinking water resources. The hidden threat to this vital ecosystem service we reveal here calls for a concerted effort to investigate the impacts of forest dieback on groundwater quality. Specifically, we identified major gaps in terms of data availability, process understanding, and interdisciplinary research that will be necessary:

### Data availability

Contrary to what one might expect, continuous data on drinking water quality, especially before purification, is difficult to obtain. While water supply utilities operationally measure water quality data, it is often not publicly accessible. European initiatives, such as the Water Framework Directive (WFD) or the annual report of the European Environmental Agency (EEA), request data to conduct large-scale assessments on groundwater quality. However, this data is often only accessible as coarsely aggregated products or with insufficient temporal coverage and metadata. The situation worsens even more if looking at other parts of the world with less dense monitoring networks or public access to data. Hence, making continuous data on groundwater quality with sufficient spatial coverage and appropriate metadata accessible is a prerequisite to catalyze research on hidden or emerging threats to drinking water quality.

Similarly, more advanced high-resolution remote sensing products are needed to disentangle different causes and types of tree mortality, such as standing deadwood, wind throw, or human-induced tree removal (Schiefer et al., 2023), and their effects on drinking water quality. Leveraging such approaches for assessments like ours, would allow capturing the effect of droughts and associated diebacks in mixed-species forests that largely do not experience complete canopy cover loss but rather scattered tree mortality. This would enable us to examine the effectiveness of silvicultural adaptation measures, such as diversifying tree species composition, that aim to alleviate or mitigate drought impacts and associated deteriorations of drinking water quality.

### Process understanding

To address the hidden threat forest dieback poses to drinking water resources, we need studies that examine the abiotic and biotic controls on nitrate leaching across the highly heterogeneous WPAs. In line with the need for advanced remote sensing products, we still lack an understanding of how the type and extent of forest dieback affect groundwater recharge and, finally, groundwater quality and how this response varies in different catchments and under different climatic conditions. From the hydrological perspective, research is needed that addresses the hydrological transport of nitrate (and potentially also other water quality parameters) in terms of concentrations and loads from the forest floor to the groundwater. This would allow us to understand N transit times (i.e., the time it takes until changes in N source availability get measurable in the groundwater) and biogeochemical retention along the flow paths.

### Interdisciplinary Collaboration

Interdisciplinary collaboration is another challenge as knowledge on data availability and process understanding exists, but it is too often tied to certain scientific disciplines, limiting our ability to identify links and potential implications. We argue that we can only understand the full dimension of the threat that forest dieback poses on drinking water quality when leveraging this knowledge in interdisciplinary research addressing the *water forest nexus* (Springgay et al., 2019). This opinion piece is such an initial effort by colleagues from hydrology, forestry, and remote sensing which should be stepped up in future collaborations. Similarly, there is an apparent lack in policy integration across the water and forest sectors (Häublein et al., 2024). Hence, policy integration is urgently needed along with the interdisciplinary research described above to address complex, cross-cutting issues arising from the increasing frequency and severity of climate extremes and their compound events (Häublein et al., 2024).

## 6 Ways forward: potential management options for safeguarding drinking water quality

To guarantee the continued provision of key ecosystem services, such as water quality protection, we need to rethink the way we conceive and manage our forests, fostering multifunctional forest landscapes that are resistant and resilient against climate extremes (Messier et al., 2019). However, there is a lack in studies analyzing the impact of the interactive effects of drought and forest management on groundwater and related drinking water quality. Studies on the impact of forest management on water quality in general suggest that options to safeguard water quality can include mixed-species forests, avoidance of clear-cuts, and the implementation of buffer stripes around surface waters (Fiquepron et al., 2013; Luke et al., 2007; Mupepele & Dormann, 2017; Osborne & Kovacic, 1993; Shah et al., 2022; Sweeney et al., 2004). Species mixing in particular may not only safeguard water quality but is also a key silvicultural adaptation strategy to climate change. This is as tree diversity facilitates forest’s buffering capacity of extreme climate events through various mechanisms. These include the insurance effect, where some species can likely compensate for the failure or reduced functioning of drought-affected species in mixtures, and the reduced competition for resources between species featuring, for instance, contrasting water-use strategies (Mahecha et al., 2024; Schnabel et al., 2021; Yachi & Loreau, 1999). Moreover, nutrient cycling can be enhanced and cycles more tightly closed in mixed-species forests due to, for instance, complementary nutrient uptake of the admixed species (Forrester, 2017), suggesting that nitrate leaching is lower in mixed compared to monospecific forests. Mixed-species forests may thus safeguard drinking water quality and could at the same time provide multiple ecosystem services in a better way than their monospecific counterparts (Messier et al. 2021). Hence, promising management strategies exist that could safeguard water quality and enhance forest stability but much more research is needed to design concrete measures such as species compositions tailored to specific site conditions, to efficiently protect our drinking water resources on the long term.

## 7 CONCLUSION

Vital forest ecosystems have been an indispensable safeguard for drinking water quality. However, the rising frequency and intensity of climate extremes, including droughts and compound events, and the associated forest dieback could jeopardize the forests’ crucial protective role. The presented proof-of-concept suggests that drinking Water Protection Areas (WPAs) are substantially affected by forest dieback and that this dieback can critically deteriorate groundwater quality and related drinking water quality. Quantifying the effects of global pulses of forest dieback on water quality and understanding the underlying drivers and controls will require more well-integrated data combined with interdisciplinary efforts for their analysis. Nevertheless, the potential scale of the threat, which we illustrated here using Germany as an example, already provides sufficient evidence to raise awareness that the world’s forests could lose their role as guardians of drinking water resources or even become a source of contamination as climate change intensifies.

## Supporting information

Supplementary Tabel S1

## Acknowledgments

We sincerely thank the Environmental Agencies of the Federal States of Germany for the provision of WPA boundaries and groundwater nitrate concentration data. We acknowledge the open provision of forest cover and forest type data by the European Environmental Agency (EEA), dominant tree-species data across Germany by the Thünen Institute, and canopy cover loss data by the Earth Observation Center (EOC) of the German Aerospace Center (DLR).

## Author contributions

CW and FS conceptualized the study. CW, FS, TK, KSt and MW designed and revised the methodology. Investigation and data curation was conducted by CW, KSz and SM. CW conducted the formal analysis and visualization. CW and FS wrote the original draft. All authors reviewed and edited the manuscript.

## Data availability

Data on forest cover and forest type was retrieved from the EEA (2018) under https://doi.org/10.2909/ab0e6d0b-699c-473d-bd5e-e5c634c8f99c. Dominant tree species were retrieved from the Thünen Insitute (Blickensdörfer et al., 2022) under https//doi.org/10.3220/DATA20221214084846. Canopy cover loss was retrieved from the EOC of the DLR under https://download.geoservice.dlr.de/TCCL/files/ (details in Thonfeld et al. 2022). Nitrate concentration data was retrieved from the Environmental Agencies of Germany’s Federal States Baden-Württemberg, Rheinland-Pfalz, Nordrhein-Westfalen, Niedersachsen and Hessen. Links to the repositories and further details are provided in the Supplementary Table S1. Boundaries of the drinking water protection areas were retrieved from Environmental Agencies of Federal States of Germany and combined into the first homogenized dataset on WPAs in Germany (data will be made openly accessible upon acceptance of this manuscript).

