## Supplementary Tabel S1 for "Forest dieback in drinking water protection areas – a hidden threat to water quality"

**Content of this file**

Table S1

**Table S1** Characteristics of selected drinking water protection areas, with >90% forest cover, >25% (A1 – A14), or <3% canopy cover loss (R1 – R6) and continuous groundwater nitrate concentrations data between 2008 and 2021. Significant breakpoints are indicated according to the year of the breakpoint and their significance level. For sites where a significant breakpoint occurred in 2018 or later, an arrow indicates the increase in the slope.  Data was retrieved from websites of the Federal States (latest access April 2024).

| Site ID | Area [ha] | Forest [%] | Broad-leaved [%] | Coniferous [%] | Forest loss [%] | Breakpoint | Source |
|---|---|---|---|---|---|---|---|
| A1 | 69 | 99,3 | 3,2 | 96,5 | 61,7 | ↑ 2020*** | https://jdkgw.lubw.baden-wuerttemberg.de |
| A2 | 2666 | 96,8 | 46,7 | 50,2 | 34,6 | 2012* | https://wasserportal.rlp-umwelt.de |
| A3 | 386 | 90,4 | 25,1 | 65,3 | 29,6 | 2013** | https://www.elwasweb.nrw.de |
| A4 | 69 | 98,7 | 8,6 | 90,5 | 43,2 | ↑ 2021*** | https://www.elwasweb.nrw.de |
| A5 | 2610 | 96,3 | 20,5 | 75,8 | 32,6 | ↑ 2018*** | http://www.wasserdaten.niedersachsen.de/cadenza/ |
| A6 | 277 | 98,2 | 27,0 | 70,1 | 26,8 | ↑ 2017*** | http://www.wasserdaten.niedersachsen.de/cadenza/ |
| A7 | 757 | 96,9 | 11,8 | 85,2 | 54,5 | ↑ 2020*** | http://www.wasserdaten.niedersachsen.de/cadenza/ |
| A8 | 491 | 98,3 | 47,5 | 50,9 | 26,5 | ↑ 2020*** | http://www.wasserdaten.niedersachsen.de/cadenza/ |
| A9 | 4841 | 97,1 | 23,0 | 73,8 | 33,8 | ↑ 2020*** | http://www.wasserdaten.niedersachsen.de/cadenza/ |
| A10 | 1352 | 99,0 | 33,1 | 66,0 | 33,4 | 2011*** | https://gruschu.hessen.de |
| A11 | 540 | 93,7 | 44,6 | 49,1 | 40,4 | - | https://gruschu.hessen.de |
| A12 | 35 | 100,0 | 36,8 | 63,4 | 35,0 | - | https://gruschu.hessen.de |
| A13 | 212 | 100,0 | 30,1 | 69,9 | 42,4 | 2013** | https://gruschu.hessen.de |
| R1 | 21 | 100,0 | 67,7 | 32,0 | 0,4 | 2015** | https://jdkgw.lubw.baden-wuerttemberg.de |
| R2 | 207 | 94,8 | 61,9 | 32,4 | 2,1 | 2011** | http://www.wasserdaten.niedersachsen.de/cadenza/ |
| R3 | 91 | 98,7 | 90,9 | 7,9 | 0,3 | 2009** | https://gruschu.hessen.de |
| R4 | 222 | 99,7 | 91,8 | 7,9 | 1,1 | - | https://gruschu.hessen.de |
| R5 | 370 | 99,1 | 12,1 | 87,1 | 0,4 | - | https://gruschu.hessen.de |
| R6 | 125 | 100,0 | 96,1 | 3,8 | 0,1 | 2016* | https://gruschu.hessen.de |
